# High-efficiency multi-site genomic editing (HEMSE) of *Pseudomonas putida* through thermoinducible ssDNA recombineering

**DOI:** 10.1101/851576

**Authors:** Tomas Aparicio, Akos Nyerges, Esteban Martínez-García, Víctor de Lorenzo

## Abstract

While single-stranded DNA recombineering is a powerful strategy for higher-scale genome editing, its application to species other than enterobacteria is typically limited by the efficiency of the recombinase and the action of native mismatch repair (MMR) systems. By building on [i] availability of the Erf-like single-stranded DNA-annealing protein Rec2, [ii] adoption of tightly-regulated thermoinducible device and [iii] transient expression of a MMR-supressing *mutL* allele, we have set up a coherent genetic platform for entering multiple changes in the chromosome of *Pseudomononas putida* with an unprecedented efficacy and reliability. The key genetic construct to this end is a broad host range plasmid encoding co-transcription of *rec2* and *P. putida*’s *mutL*_E36K_^PP^ at high levels under the control of the P_*L*_/*c*I857 system. Cycles of short thermal shifts of *P. putida* cells followed by transformation with a suite of mutagenic oligos delivered different types of high-fidelity genomic changes at frequencies up to 10% per single change—that can be handled without selection. The same approach was instrumental to super-diversify short chromosomal portions for creating libraries of functional genomic segments—as shown in this case with ribosomal binding sites. These results enable the multiplexing of genome engineering of *P. putida*, as required for metabolic engineering of this important biotechnological chassis.

## INTRODUCTION

DNA recombineering was first developed in the early 2000s (Datsenko and Wanner, 2000; Yu et al., 2000) as a genetic technology for replacing genomic segments of *E. coli* with synthetic double stranded (ds) DNA by means of the DNA exchange mechanism brought about by the Red system of phage lambda. While the native approach involves 3 proteins (a β-recombinase, an exonuclease and the γ protein which protects free ds-ends of DNA from degradation by RecBCD), it turned out that the Red-β protein sufficed to promote invasion of the replication fork by single-stranded oligonucleotides incorporated as Okazaki fragments (Ellis et al., 2001). If such oligonucleotides were designed to carry mutations, the resulting changes could be inherited at considerable frequencies upon subsequent rounds of DNA segregation. The key value of this approach is that by using cocktails of mutagenic oligonucleotides and either manual or automated cycles of Red expression/oligonucleotide transformation one can enter simultaneous changes in many genomic sites and/or saturate given DNA stretches with specific or random mutations (Wang et al., 2009; Nyerges et al., 2016; Nyerges et al., 2018). These methods have been improved further by using host strains transiently disabled in mismatch repair and by enriching mutants through Cas9/gRNA-based counterselection of wild-type sequences (Costantino and Court, 2003; Jiang et al., 2013; Nyerges et al., 2014; Nyerges et al., 2016; Ronda et al., 2016; Oesterle et al., 2017). While these technologies work well in *E. coli*, they are difficult to transplant directly to non-enteric bacteria. Yet, their applicability to species such as *Pseudomonas putida* has a special interest because of the value of environmental microorganisms as useful platforms for metabolic engineering (Nikel et al., 2014; Nikel et al., 2016; Martínez-García and de Lorenzo, 2019). Attempts of functional expression of the lambda Red system in various species of *Pseudomonas* have been reported, but recombination frequencies were low in the absence of selection (Lesic and Rahme, 2008; Liang and Liu, 2010; Luo et al., 2016; Chen et al., 2018; Yin et al., 2019). Red-like counterparts found in *Pseudomonas* prophages have been more successful to the same ends. For example, the RecET recombinase/exonuclease pair of *P. syringae* has been instrumental for executing a suite of manipulations in this species (Swingle et al., 2010a; Bao et al., 2012). Furthermore, bioinformatic mining of *Pseudomona*s-borne recombinases from known protein families (i.e., Redβ, ERF, GP2.5, SAK, and SAK4; Lopes et al., 2010) followed by experimental validation of the most promising in a standardized recombineering test exposed two new enzymes (Ssr and Rec2: Aparicio et al., 2016; Ricaurte et al., 2018; Aparicio et al., 2020). These recombinases delivered a comparatively high level of activity in the reference strains *P. putida* KT2440 and its genome-reduced derivative *P. putida* EM42. Still, numbers were way below those reported for *E. coli*. Furthermore, the action of the endogenous MMR system of this bacterium impeded single nucleotide changes (i.e., A to T, mismatch A:A) that were efficiently fixed by the indigenous *mutS/mutL* device (Aparicio et al., 2016; Aparicio et al., 2019b).

In this work we have set out to overcome the bottlenecks to efficacious recombineering in *P. putida* mentioned above. The approach builds on the apparently superior ability of the Rec2 recombinase to promote DNA annealing with exogenous synthetic oligonucleotides during chromosomal replication. By playing with a stringent expression system for *rec2*, applying multiple cycles of recombinase production/ oligonucleotide transformation and reversibly inhibiting the MMR system during a limited time window we report below high-fidelity recombination frequencies that approach those achieved with the archetypal Red-based system (Datsenko and Wanner, 2000). This opens genome-editing possibilities in this environmental bacterium that were thus far limited to strains of *E. coli*, closely related enteric species (Nyerges et al., 2018; Szili et al., 2019) and some lactic acid bacteria (van Pijkeren et al., 2012).

## RESULTS AND DISCUSSION

### Optimization of Rec2 and MutL_E36K_^PP^ delivery for ssDNA recombineering

The bicistronic gene cassette of pSEVA2514-*rec2*-*mutL*_E36K_^PP^ (Fig. 1) was developed earlier for examining the hierarchy of recognition of different types of single nucleotide mispairs by the native mismatch repair (MMR) system of *P. putida* (Aparicio et al., 2019b). During the course of that work, we noticed that a short, transient, thermal induction of the Rec2 recombinase increased very significantly ssDNA recombineering (~ 1 order of magnitude) as compared to the same with an expression device responsive to 3-methyl-benzoate (i.e. *xylS*/P*m*). Although the reason for this improvement is not entirely clear, it may have resulted from [i] the short-lived, high-level transcription of the otherwise toxic recombinase—as compared to the permanent hyperexpression caused by the chemically-inducible system, [ii] thermal inactivation of ssDNA nucleases and thus improved survival of the mutagenic oligonucleotides *in vivo* or [iii] a combination of both. In any case, the average frequency of single-base replacements in just one single-shot recombineering test was in the range of 1E^-2^ mutants per viable cell. This was high compared to previous recombineering efforts in this bacterium (Aparicio et al., 2016) but still low for identifying mutations without a selectable phenotype. We however speculated that by multi-cycling the procedure with short thermal pulses of induction/resetting of recombinase expression and transformation with mutagenic oligos, such frequencies could be added at each cycle, eventually resulting in high nucleotide replacement rates. A second realization of (Aparicio et al., 2019b) was that transient co-expression of the dominant allele MutL_E36K_^PP^ of the MMR system of *P. putida* along with the *rec2* gene in plasmid pSEVA2514-*rec2*-*mutL*_E36K_^PP^ (Fig. 1) virtually eliminated recognition of any type of base mispairings in DNA. This allowed entering all classes of nucleotide replacements which would otherwise be conditioned by MMR—without triggering a general mutagenic regime. Yet, note that both activities (Rec2 and MutL_E36K_^PP^) were delivered *in vivo* with a high copy number vector with an origin or replication (RSF1010) of unknown thermal sensitivity. This may result in some instability upon thermal cycling of the procedure for boosting recombineering efficiency (see below). In order to determine the best plasmid frame for *rec2*-*mutL*_E36K_^PP^ transient expression, the cognate DNA segment was recloned in vectors pSEVA2214 (RK2 origin or replication, low copy number) and pSEVA2314 (pBBR1 origin, medium copy number) as shown in Fig. 1A. Recombineering tests were then carried out with oligonucleotide NR, which generated a double mutation in *gyrA* endowing resistance to nalidixic acid (Nal^R^) by means of two MMR-sensitive changes G ➔ A and C ➔ T. In parallel, another MMR-insensitive change A ➔ C was also tested with oligonucleotide SR that mutated *rpsL* for making cells resistant to streptomycin (Sm^R^). The results of this test indicated pSEVA2314-*rec2*-*mutL*_E36K_^PP^ as the preferred construct of reference for the muti-site mutagenesis platform presented below. On the basis of the above we set out to recreate in *P. putida* the same conditions that enabled implementation in *E. coli* of high-efficiency ssDNA recombineering protocols such as MAGE (Wang et al., 2009), DIvERGE, (Nyerges et al., 2018), and pORTMAGE (Nyerges et al., 2016)—and thus expand frontline genomic editing methods towards that environmentally and industrially important bacterium.

**Figure 1.**
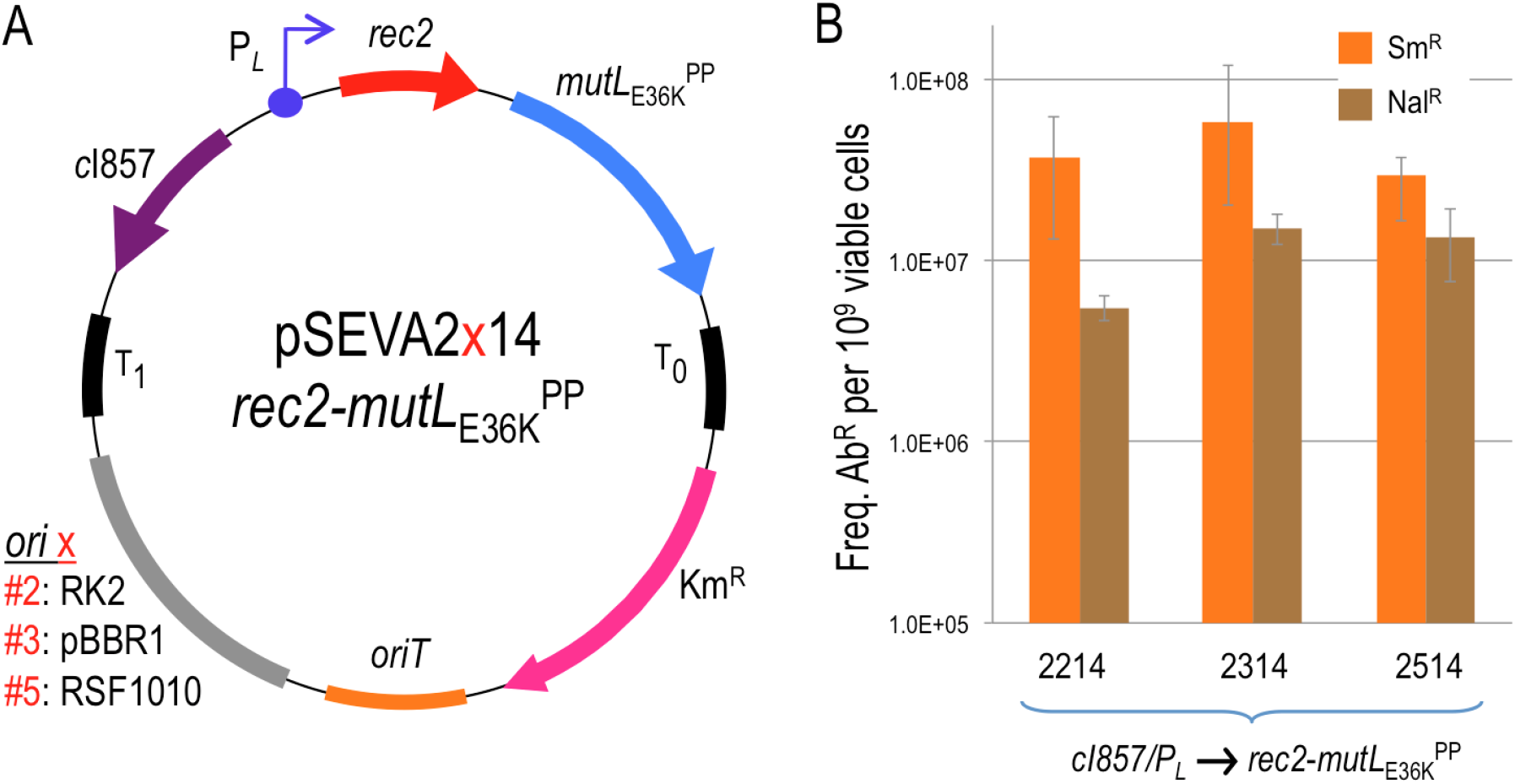
Influence of plasmid copy number in the editing efficiency of the heat-induced *rec2-mutL*_E36K_^PP^ genes. **A)** Genetic map and structure of plasmids used in this study. The figure shows the plasmids tested, all having the same elements with the exception of the origin of replication, represented with “x”. T_0_ and T_1_, transcriptional terminators; Km, Kanamycin resistance gene; *ori*T, origin of transfer; *c*I857-P_*L*_, temperature inducible expression system; *rec2*, recombinase; *mutL*_E36K_^PP^, dominant-negative allele of *mutL*; *ori* x (origin of replication): #2, RK2 (low copy number); #3, pBBR1 (medium copy number); #5, RSF1010 (medium-high copy number). Pictures are nor drawn to scale. **B)** Recombineering assays with *P. putida* EM42: the strain harbouring each pSEVA2×14-*rec2*-*mutL*_E36K_^PP^ variant was subjected to recombineering with oligos SR and NR upon heat induction of the *c*I857-P_*L*_ expression system. After overnight recovery, culture dilutions were plated on LB-Sm (SR oligo) and LB-Nal (NR oligo) to estimate the number of allelic changes. Culture dilutions plated on LB allowed viable cells counting. Column values represent mean recombineering frequencies (mutants per 10^9^ viable cells) of two independent experiments with the standard deviation.

### Cyclic pulses of rec2/mutL*_E36K_^PP^* expression enable a high level of single-nucleotide substitutions

The first issue at stake was determining the frequencies of mutations caused by using a cocktail of oligonucleotides targeting 5 genes representative of diverse genomic locations, different types of nucleotide changes and associated or not to selectable traits upon multiple ssDNA recombineering cycles. The genes at stake, their position in the chromosomal map, the cognate phenotypes and the type of replacements brought about by the corresponding mutagenic ssDNAs are summarized in Fig. 2. They were all designed to pair sequences in the lagging strand of the replication fork in each of the replichores of the *P. putida* genome according to (Aparicio et al., 2016). Note that the experiments were run with *P. putida* EM42, not with the archetypal strain KT2440. This is because it is a recA^+^ derivative of the EM383 genome-streamlined variant that has higher endogenous levels of ATP and NAD(P)H and has thus become a preferred metabolic engineering platform (Martinez-Garcia et al., 2014). Moreover, the modifications entered in *P putida* EM42 make this strain more tolerant to pulses of high-temperature (Aparicio et al., 2019a), as repeatedly applied throughout this work (see below). The cyclic recombineering protocol (see Transparent Methods for details) is summarized in Fig. 3, and it basically involves four steps: [i] growing cells, [ii] triggering thermal induction of *rec2* and *mutL*_E36K_^PP^ genes by a short heat-shock, [iii] preparing competent cells for electroporation with the mutagenic oligonucleotides and [iv] recovering the culture for a new cycle.

**Figure 2.**
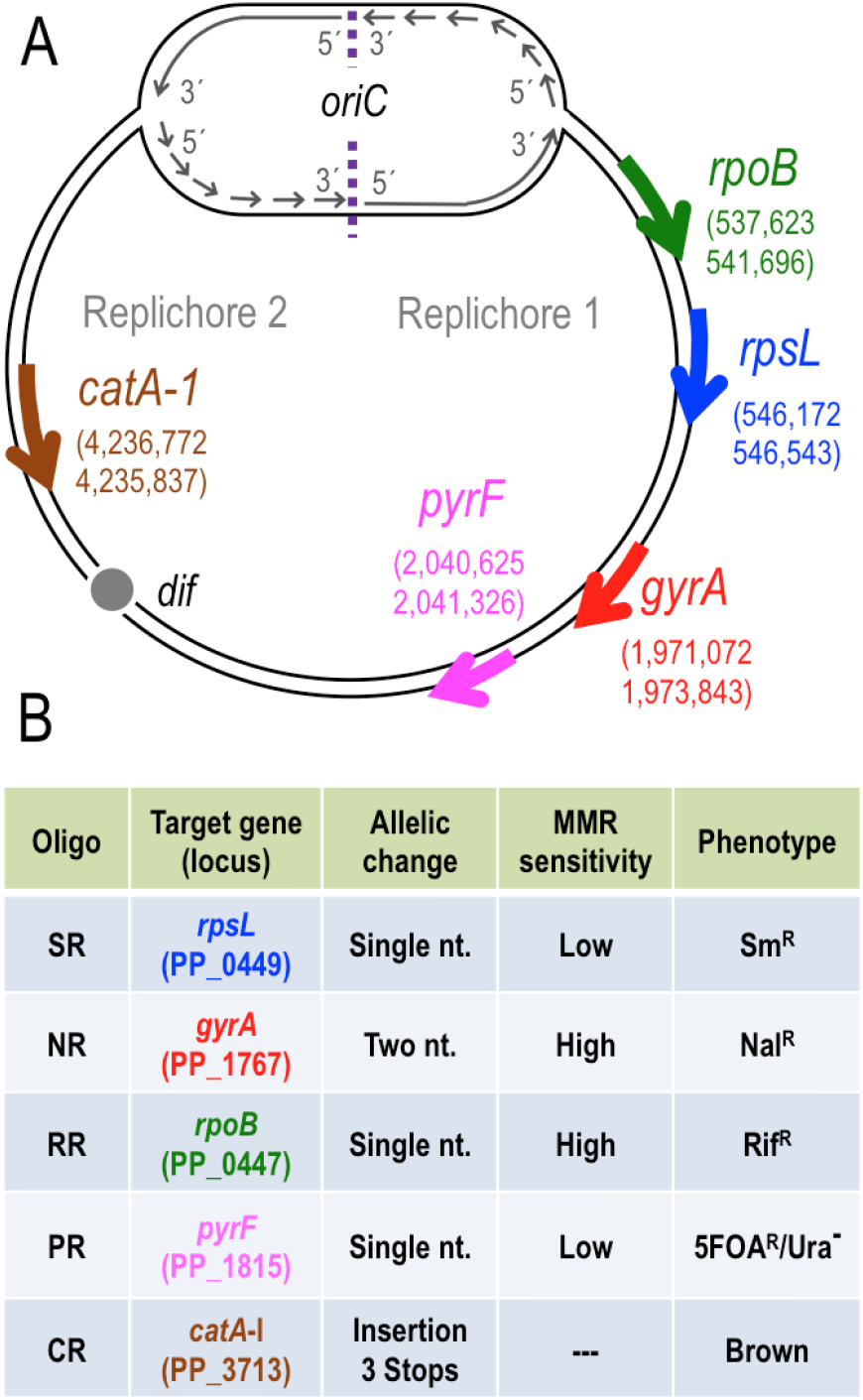
Target genes and recombineering oligonucleotides used for HEMSE. **A)** The 5 genes selected as targets for recombineering are represented in the chromosomal map of *P. putida* KT2440 with gene coordinates and strand orientation. *oriC* and *dif* regions are shown to define the two replichores in the genomic map. Pictures are not drawn to scale. **B)** The main features of recombineering oligonucleotides used to assay HEMSE are shown: name of oligo, target gene with its locus tag, type of allelic replacement, level of Mismatch Repair (MMR) sensitivity of the allelic changes and the cognate phenotypes produced. See Supplementary Tables S1 and S3 for additional information.

**Figure 3.**
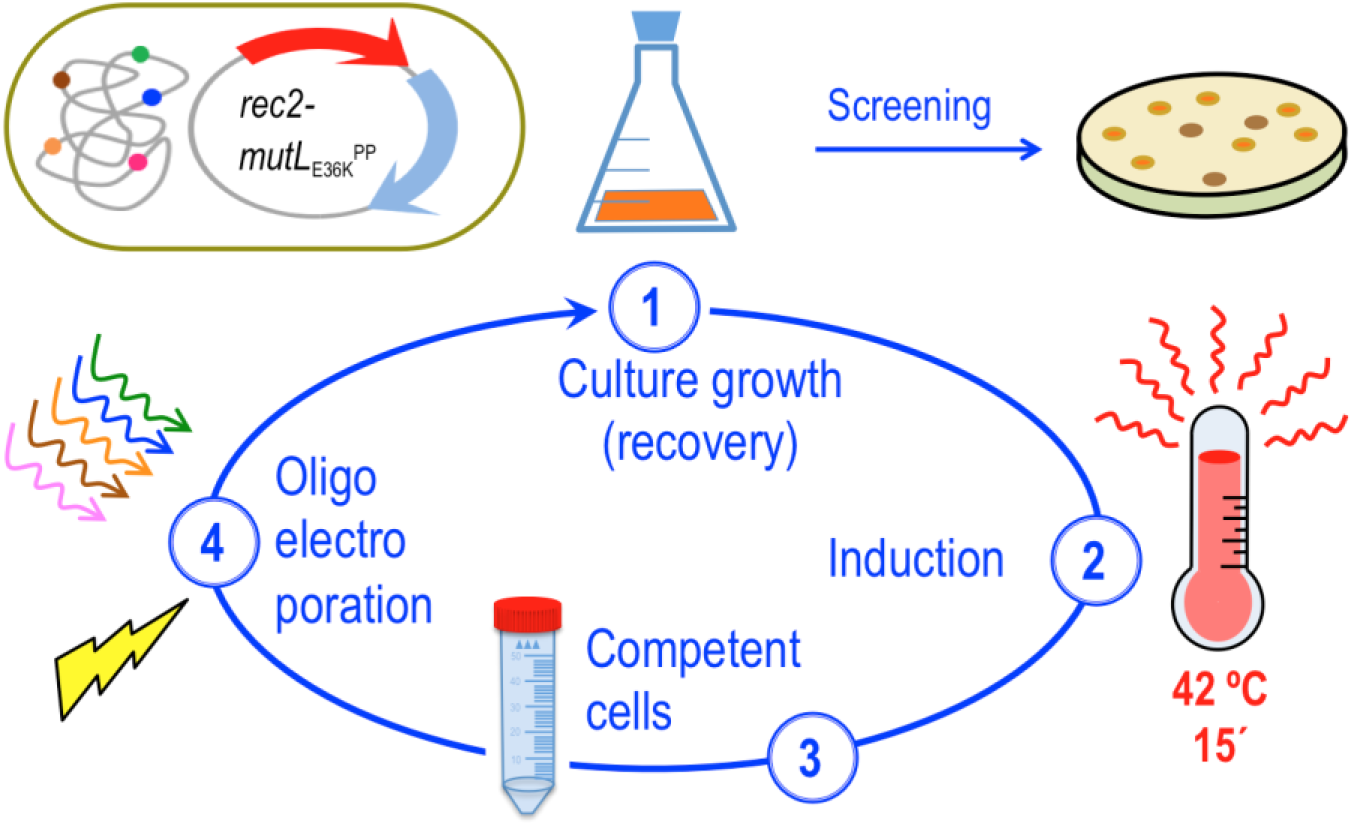
Scheme of HEMSE cycle. The main steps of the procedure are depicted: cultures of *P. putida* EM42 (pSEVA2314-*rec2*-*mutL*_E36K_^PP^) grown at OD_600_ = 1.0 are induced by a heat-shock at 42 °C/ 15 minutes, then competent cells are prepared and transformed with recombineering oligonucleotides. After recovery on fresh media at 30 °C/ 170 rpm, cultures enter in the next round of HEMSE by applying the induction step. Screening of allelic replacements within a given cycle is performed after recovery by plating culture dilutions on the appropriate solid media (see Transparent Methods for details)

The results of applying multiple recombineering cycles to *P. putida* EM42 (pSEVA2314-*rec2*-*mutL*_E36K_^PP^) with the oligonucleotides listed in Fig. 2B are shown in Fig. 4A. Note that the frequencies of mutant appearance increased during the runs from 2.8E^-3^ (1 cycle) ➔ 9.3E^-2^ (10 cycles) in the case of *gyrA*, to 4.8E^-2^ ➔ 2.0E^-1^ for *pyrF* under the same conditions. In the best-case scenario (i.e. gene *rpsL*), the frequencies multiplied by 24-fold, reaching a remarkable 21%. After the 10th cycle, these figures are thus close to the rates reported in *E. coli* with the archetypal Red-β system of phage lambda and also to the theoretical limit of recombineering frequencies (25 %) that stems from segregation of one allelic change after two rounds of genome replication (Wang et al., 2009; Nyerges et al., 2016). It is worth to mention that control strain *P. putida* EM42 harbouring insert less vector pSEVA2314—but transformed with the same mutagenic oligonucleotides—gave rise to recombineering frequencies ~ 1E^-5^/1E^-6^ mutants/viable cells per cycle for single changes (Fig. S1). Given that these background levels are higher with thermal induction than with chemical induction (Ricaurte et al., 2018), it is plausible that heat shock intrinsically improves recombineering regardless of the action of heterologous recombinases. As a matter of fact, purely endogenous ssDNA recombineering at significant frequencies has been reported in a variety of Gram-negative bacteria, including *E. coli* and *Pseudomonas syringae* (Swingle et al., 2010b), a fact that play in our favour for establishing the methodology in *P. putida*.

**Figure 4.**
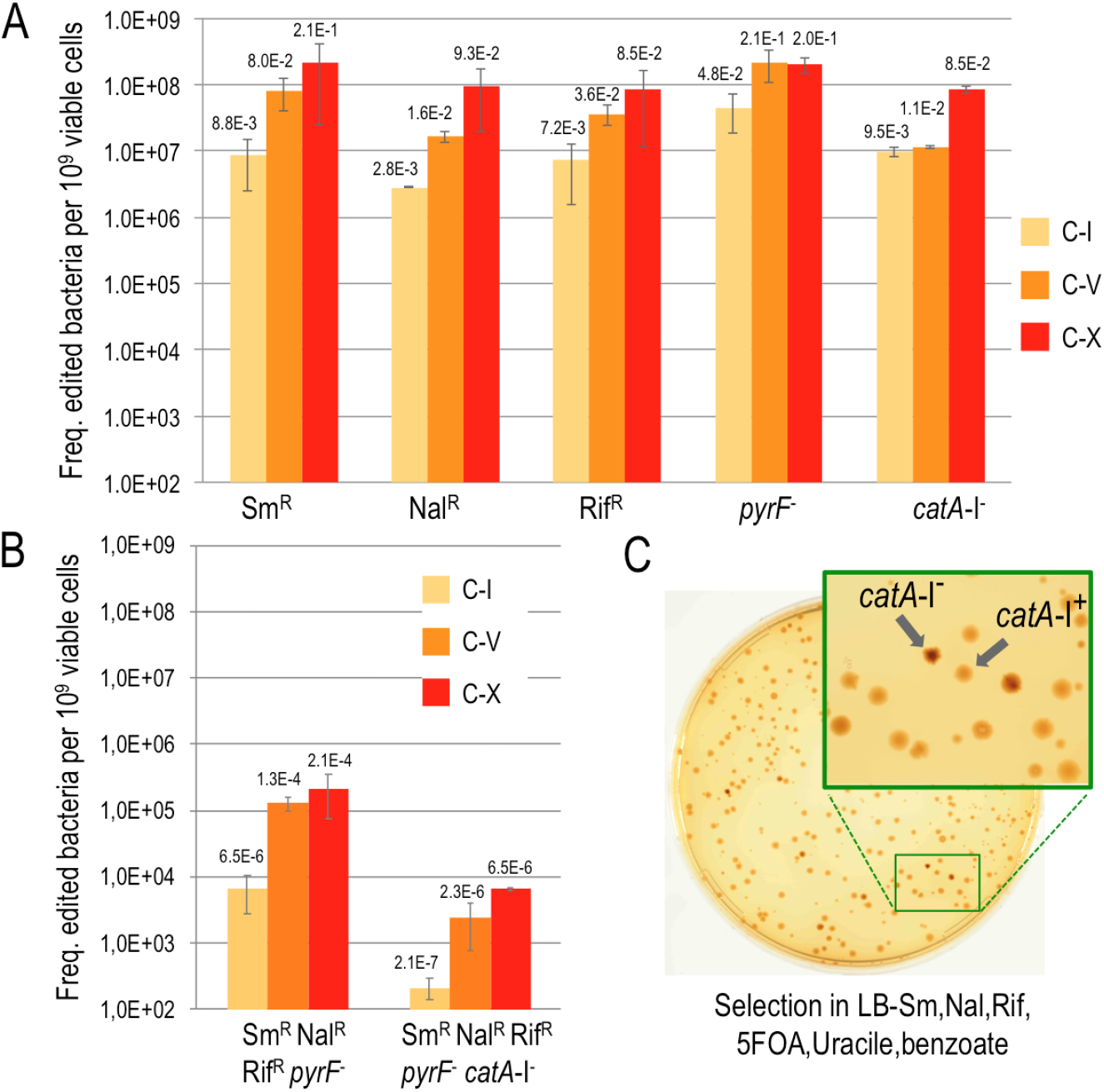
Editing efficiencies HEMSE. **A)** Editing efficiencies of single changes were assayed applying 10 cycles of HEMSE to *P. putida* EM42 (pSEVA2314-*rec2*-*mutL*_E36K_^PP^) using an equimolar mixture of oligos SR, NR, RR, PR and CR. After recovery steps of cycles n° 1 (C-I), n° 5 (C-V) and n° 10 (C-X), appropriate dilutions of the cultures were plated on LB to estimate viable cells and also on LB solid media supplemented with Sm, Nal, Rif, 5FOA-Ura or benzoate to enumerate allelic replacements. Colonies growing on Sm, Nal, Rif or 5FOA-Ura were counted as allelic changes, while brown - catechol accumulating-colonies growing in LB-benzoate were counted as *catA*-I^-^ clones. Recombineering frequencies of single replacements at each cycle were normalized to 10^9^ viable cells and the medias of two independent replicas were plotted with standard deviations. Absolute recombineering frequencies (mutants per viable cell) are also depicted over the bars. **B)** From the same experiments explained above, editing efficiencies of multiple changes were analysed. Dilutions of C-I, C-V and C-X were plated on LB-SmNalRif-5FOA-Ura and LB-SmNalRif-5FOA-Ura-benzoate solid media, allowing the estimation of, respectively, quadruple (Sm^R^ Nal^R^ Rif^R^ *pyrF*) and quintuple (Sm^R^ Nal^R^ Rif^R^ *pyrF*^-^ *catA*-I^-^) editions. Results were represented as in A). **C)** A representative plate of quintuple screening at C-X is shown. The zoom-up shows colonies with the characteristic dark-brown phenotype of *catA*-I^-^ clones.

The most remarkable outcome of the operations shown in Fig. 4A was that such high figures enabled manual screening of inconspicuous mutations, thus avoiding the need of adding a genetic counterselection device (e.g., CRISPR/Cas9) for identifying rare changes. Since these results accredited the value of multi-cycling thermoinduction of the bicistronic *rec2*-*mutL*_E36K_^PP^ operon of pSEVA2314-*rec2*-*mutL*_E36K_^PP^ for raising ssDNA recombineering efficiency, the next obvious question was whether the high figures could afford simultaneous multi-site genomic editing with mixtures of mutagenic oligos, in a fashion reminiscent of the MAGE (Multiplex Automated Genome Engineering) process available for *E. coli*.

### Multi-site editing of non-adjacent genomic locations

Given average individual mutation rates of 10% after 10 thermal recombineering cycles and assuming they are separately maintained when cells face a cocktail of mutagenic oligonucleotides one can predict frequencies of 1% double changes all the way to 0.001 % mutants (1E^-5^) of genomes with all the 5 changes in the absence of any phenotypic advantage. To test this prediction we subjected a culture o *P. putida* EM42 (pSEVA2314-*rec2*-*mutL*_E36K_^PP^) to 10 cycles of thermoinduced recombineering (see Transparent Methods) with re-transformation in each cycle with an equimolar mixture of oligos SR, NR, RR, PR and CR (Fig. 2B; Supplementary Table S1) so that all possible changes could be entered in the same cells. Emergence of multiple (i.e. quadruple and quintuple) mutations in the population was then monitored at cycles I, V and X and their frequencies recorded. Fig. 4B shows the results of such a procedure. The data exposed a good match between the theoretical expectation of multiple changes and the actual figures, although the evolution of the mutation rates was not linear. At cycle #1, single changes showed recombineering frequencies barely below 1E^-2^ mutants per viable cell. If we take that as a reference, theoretical frequencies acquisition of 4 and 5 changes would be 1E^-8^ and 1E^-10^ respectively, while actual numbers were way higher (6E^-6^ and 2E^-7^). By cycle #5, single changes reached average recombineering above 5E^-2^. The gross theoretical prediction for simultaneous appearance of 4 and 5 changes would be as low as 6E^-6^ and 3E^-7^. Yet, again, the actual experiments yield 1.3E^-4^ and 2.3E^-6^ mutants per viable cell for quadruple and quintuple mutants. By cycle #10, however, the scenario was different. Single changes appeared at frequencies ~ 1E^-1^, close to the theoretical maximum of recombineering efficiency (2.5 E^-1^). In this case, predicted frequencies for 4-5 changes ranged 1E^-4^ and 1E^-5^, which were very similar to the actual numbers delivered by the experiment i.e. 2E^-4^ and 6E^-6^.

The results above suggested that, during the first 5 recombineering cycles a strong co-selection phenomenon occurs. Appearance of multiple mutations fall 2-3 logs higher than expected, suggesting that cells undergoing ssDNA incorporation in specific loci are more prone to incorporate changes in other genomic locations. This phenomenon, which has been observed before (Carr et al., 2012; Gallagher et al., 2014) could be due to differences in the ability of single cells in a population to uptake exogenous ssDNA upon electroporation. Regardless of the specific mechanisms, the results of Fig. 4B show that multi-cycle recombineering boosts mutagenic frequencies through single to quintuple changes. Yet, while 10 cycles appear to reach saturation at single sites, it is plausible that additional runs could enrich further the population in multi-edited bacterial cells. Taken together, the experiments of Fig. 4 document the power of the hereby described method for simultaneously targeting 5 genomic sites of *P. putida* for desired mutations. On this basis we propose to call the entire workflow High-Efficiency Multi-site Genomic Editing (or HEMSE). The method is conceptually comparable to Multiple Automated Genome Editing developed for *E. coli* (MAGE, (Wang et al., 2009)) but it lacks (thus far) the automation aspect.

Since a growing culture of *P. putida* in LB typically ranges 10^8^-10^9^ cells/ml from early exponential to early stationary phase, we speculated that the maximum number of genes that could be edited in an HEMSE experiment of this sort with mixed oligos in the absence of any selective advantage or phenotypic screening could be ~ 8-9. This is clearly not enough for massive changes of the sort necessary for e.g. recoding a whole genome (Isaacs et al., 2011) or reassigning/erasing specific triplets (Ostrov et al., 2016). Fortunately, in most typical metabolic engineering endeavours, the issue is not so much entering many defined mutations in given chromosomal sites but fostering the system to explore a solution space by letting it come up with many combinations—the most successful of which can be enriched and subject to further mutation rounds. This effect can be exacerbated if the mutagenic oligos boost the diversification of e.g. regulatory sequences, so their combination generates fluctuations in the stoichiometry of a multi-gene pathway (Hueso-Gil et al., 2019)—or they create variants of the same protein with different activities by diversifying specific segments. The technical issue shared by all these scenarios is the focusing of the diversification in a defined sequence window of the genomic DNA. In this context the question is whether the above described HEMSE is instrumental to this end also—as the recombineering-based method to the same end called DIvERGE is in *E. coli* and related enterobacteria (Nyerges et al., 2018).

### Diversification of the SD motif context creates new functional RBSs in P. putida

In order to have a tractable proxy of generation *in vivo* of large libraries of functional DNA sequences in the *P. putida* genome, the experimental setup shown in Fig. 5 was developed. In it, a Tn*7* mini-transposon vector was inserted with the *gfp* gene downstream of the constitutive promoter P_*EM7*_ but lacking a recognizable Shine Dalgarno (SD) sequence for translation initiation. The hybrid transposon was subsequently inserted in the cognate *att*Tn*7* site of the *P. putida* EM42 chromosome (see Transparent Methods) from which it was expectedly unable to produce any detectable fluorescence. The resulting strain (*P. putida* TA245, Supplementary Table S2) was transformed with pSEVA2314-*rec2*-*mutL*_E36K_^PP^ and used in recombineering experiments with oligonucleotides designed for creating RBS variants. The business parts of such oligonucleotides are shown in Fig. 5B. As controls we used oligos named RBS-C_6_ and RBS-C_9_. These ssDNA enter respectively a short or an extended Shine-Dalgarno (SD) sequence, 8 bp upstream of the start codon of the *gfp* gene using as a reference the *P. putida* 16S ribosomal gene an containing the core for optimal translation 5’-GAGG-3’ (Shine and Dalgarno, 1975; Kozak, 1983; Farasat et al., 2014). For RBS diversification we used oligonucleotides RBS-Deg_6 and_ RBS-Deg_9_ (Fig. 5B), which include soft randomized sequences with discrete changes R (A or G) that cover 6 degenerated positions with a potential to generate 64 (= 2^6^) combinations. This was expected to create a large population of RBS of different efficacies, which could be quantified through fluorescent emission of individual cells.

**Figure 5.**
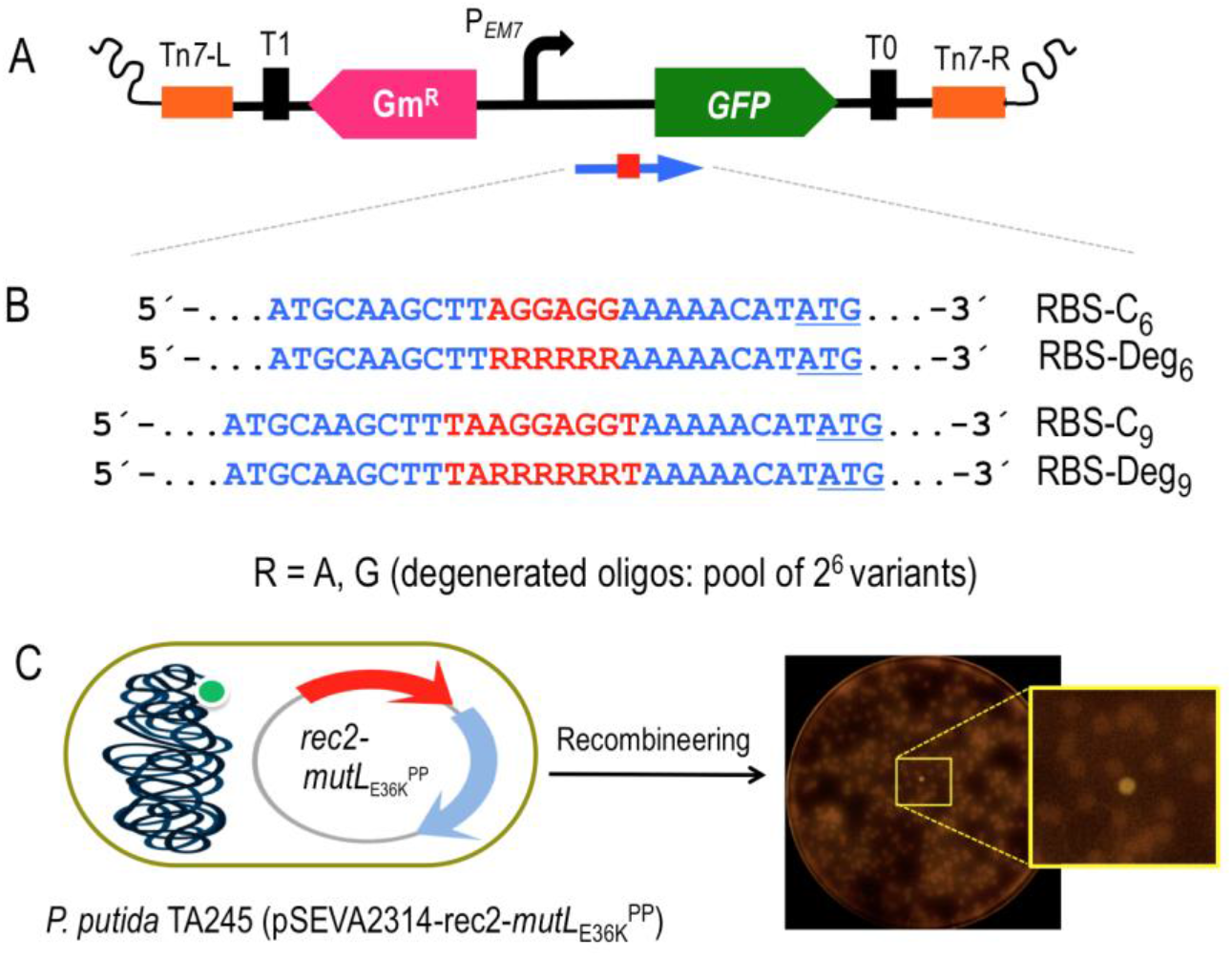
Diversification of the *gfp* Shine Dalgarno motif. **A)** A mini-Tn*7* transposon bearing the *gfp* gene devoid of its original SD sequence and under the control of the constitutive P_*EM7*_ promoter was constructed. The elements depicted are: Tn*7*-L and Tn*7*-R, left and right Tn*7* sites; T_0_ and T_1_, transcriptional terminators; Gm^R^, Gentamicin resistance gene; P_*EM7*_, constitutive promoter; GFP, Green Fluorescent Protein gene. A blue arrow represents the target region of recombineering oligonucleotides aimed to reconstruct the *gfp* ribosome binding site (shown as a red square). **B)** Partial sequence of the four recombineering oligonucleotides (Supplementary Table S1) designed to introduce SD motifs upstream the *gfp* gene. RBS-C_6_ and RBS-C_9_ insert, respectively, the semi-canonical AGGAGG and the canonical TAAGGAGGT SD motifs eight nucleotides upstream the ATG start codon of the *gfp* (underlined). RBS-Deg_6_ and RBS-Deg_9_ insert the randomized sequences RRRRRR and TARRRRRRT, where R stands for A (adenine) or G (guanine). Each degenerated oligonucleotide comprises a pool of 64 variants (2^6^) with all possible combinations A/G. **C)** The mini-Tn*7* device was inserted in the *att*Tn*7* site of *P. putida* EM42. Upon transformation with pSEVA2314-*rec2*-*mutL*_E36K_^PP^, the resulting strain *P. putida* TA245 (pSEVA2314-*rec2*-*mutL*_E36K_^PP^) was subjected to one HEMSE cycle with the recombineering oligos in independent experiments. After plating in LB-GmKm-charcoal, GFP positive clones were identified. A plate from the screening of RBS-Deg_9_ is also shown, with a magnification of a fluorescent colony.

For the experiments described below, *P. putida* TA245 (pSEVA2314-*rec2*-*mutL*_E36K_^PP^) was separately subject to one recombineering cycle with each of four oligos of Fig. 5B, after which cells were diluted and plated in charcoal-LB agar for easing visual detection of colonies emitting low fluorescence on a black background. Positive controls RBS-C_6_ and RBS-C_9_ allowed the estimation of editing frequencies as GFP^+^ cells/total number of cells, which resulted in 5.9E^-4^ and 9.9E^-4^, respectively. Those values were relatively low as compared to the recombineering efficiencies reported above for single changes (~ 1E^-2^). This could be possibly due to the shorter homology arms the oligos (30 nt) and the extended sequence inserted between them (Aparicio et al., 2020). Yet, these figures provided a reference for subsequent quantification of the effect of soft-randomized oligos RBS-Deg_6 and_ RBS-Deg_9_. After treatment with these last, cultures were diluted and plated for inspection of ~ 9000 colonies resulting of each recombineering experiment. Visual screening of the colonies revealed the appearance of 67 and 53 fluorescent clones coming, respectively, from experiments with RBS-Deg_6_ and RBS-Deg_9_. These 120 clones were picked up for further analysis. PCR and sequencing of the region upstream the *gfp* gene allowed identification of 14 variants of RBS-Deg_6_ and 17 variants of RBS-Deg_9_, the GFP levels of which were measured by flow cytometry. The results plotted in Fig. 6 show that the different variants delivered emissions ranging from very low to high fluorescent levels across a 20-fold change span. It is worth to highlight that the best RBS of the series (Strain Code #33, Fig. 6) has a perfect match with the complementary sequence of the last 9 nt of the 16S ribosomal RNA of *P. putida* (PP_16SA). Other clones (e.g. #32; RBS = 5’-AAGGAG-3’) displayed also high fluorescence levels. Interestingly, a few productive variants contained the same 6-nt sequence in the degenerated region regardless of the type of randomized oligo (e.g. #32 = #30, #26 = #16, #14 = #11 and #23 = #10). While most of the high-signal variants belong to the longer RBS-C_9_-borne clones, the comparison of signals does not support the hypothesis that longer complementarity to the 16s ribosomal sequence correlates with more efficient translation. Other factors have been proposed to affect translation efficacy of RBS variants, such as the stability and secondary structure of RNA and transcriptional efficiency (Chen et al., 1994; Salis et al., 2009). Regardless of the possible biological significance of the results, the data of Fig. 6 certifies the efficacy of the HEMSE platform to generate diversity in specific genomic segments—a welcome feature which can doubtless be multiplexed to other chromosomal locations as required.

**Figure 6.**
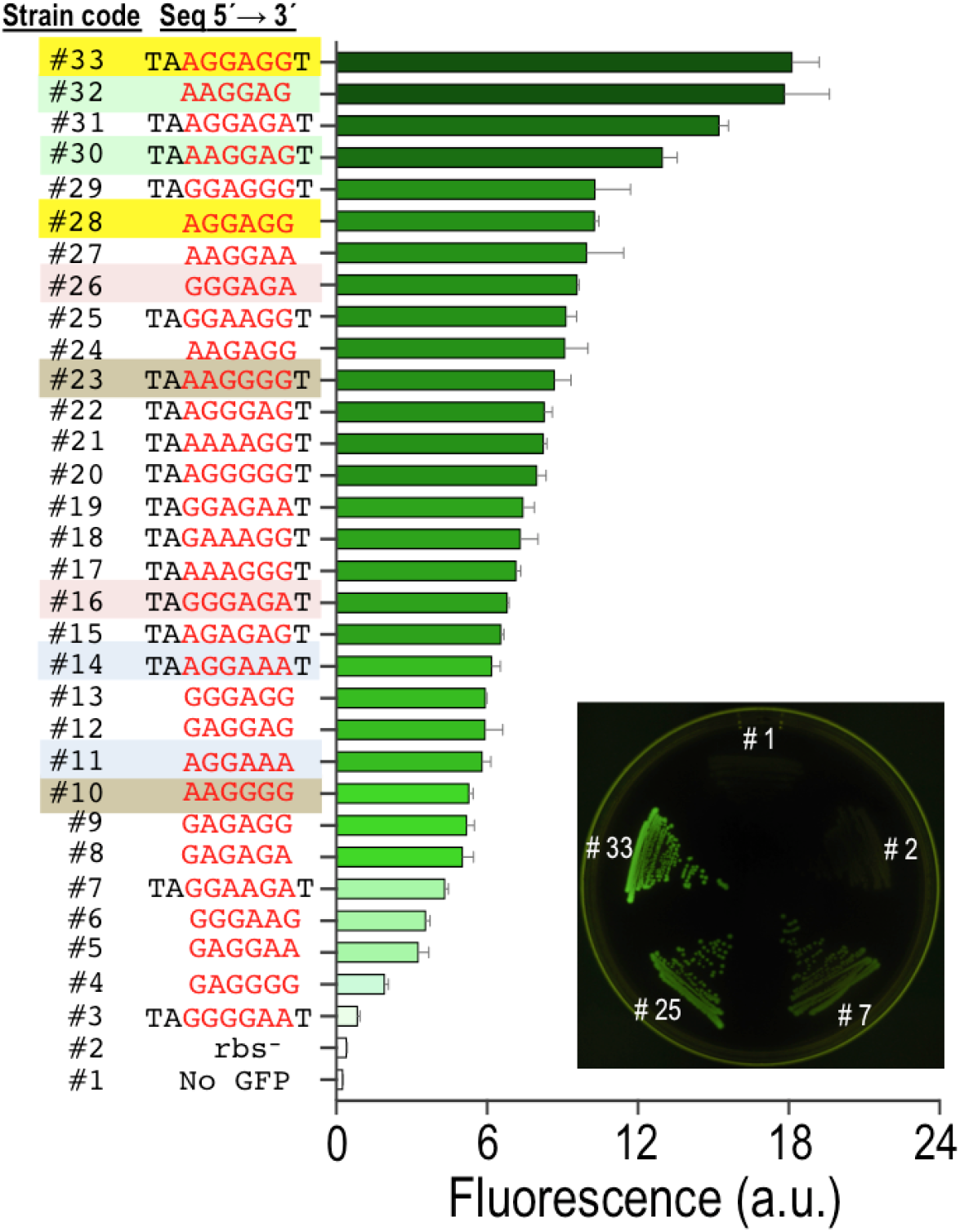
Characterization of the diversified library of SD sequences. The screening of the HEMSE experiments performed in *P. putida* TA245 (pSEVA2314-*rec2*-*mutL*_E36K_^PP^) with RBS-Deg_6_ and RBS-Deg_9_ yielded 31 variants in the ribosome binding site of the *gfp* gene. The GFP expression of this library was analysed by flow cytometry including two negative controls: *P. putida* EM42 (Strain #1), in which there is no *gfp* gene, and also the ancestral strain *P. putid*a TA245 (pSEVA2314-*rec2*-*mutL*_E36K_^PP^), harbouring the mini-Tn*7* and the *gfp* gene but lacking the original SD sequence (Strain #2). The plot shows the mean fluorescent emission of individual clones from two biological replicas with standard deviations. A Strain Code was assigned to each variant analysed and the putative SD sequence identified 8 nt upstream the *gfp* start codon is shown. Variants showing identical sequence in the randomized region are highlighted with the same color. A plate of LB-charcoal agar with streaks of the two controls and three representative clones exhibiting low-, medium and high signal rates (#7, #25 and #33, respectively) is also depicted under UV light.

### Conclusion

In this work we have merged and adapted to *P. putida* and in a single platform 3 of the most efficacious genetic tools available to metabolic engineers for generating diversity *in vivo* focused into a predetermined number of chromosomal DNA segments: ssDNA recombineering (Wang et al., 2009), portable MAGE (Nyerges et al., 2016) and DIvERGE (Nyerges et al., 2018). Although conceptually identical to such methods already applied to *E. coli*, their recreation in a non-enterobacterial species involved [i] the search and testing of functional equivalents of the parts involved but recruited from *Pseudomonas* genomes and [ii] adaptation and optimization to the distinct physiology of the species and strain at stake. While we have not made a side-by-side comparison of the frequencies resulting from standard MAGE in *E. coli* and the ones presented in this work, numbers in the range of 10% replacements after 10 recombineering cycles could be sufficient to implement the same powerful method in *P. putida*. We are reluctant, however, to use the same acronym, because the automation feature is not in sight and the multiplexing still problematic with the current efficiencies.

One can envision various ways through which HEMSE could be further improved. End-terminal degradation of the mutagenic oligos *in vivo* does not seem to be an issue: performance of 5’-phosphorothioated ssDNA (which cannot be degraded by exonucleases (Wang et al., 2009) is indistinguishable from non-phosphorothioated equivalents (Aparicio et al., 2020). However, the nature and origin of the recombinase that catalyzes invasion of the DNA replication fork by the synthetic oligo makes a considerable difference (Chang et al., 2019). It is possible that such recombinases act in concert with additional endogenous proteins that could be characteristic of each species (Caldwell et al., 2019; Yin et al., 2019). It seems thus desirable that future alternatives to the Rec2 activity encoded in pSEVA2314-*rec2*-*mutL*_E36K_^PP^ (Fig. 1) are mined in *Pseudomonas* genomes and phages—by themselves or in combination with other complementary genes. It should be straightforward to then replace the *rec2* of pSEVA2314-*rec2*-*mutL*_E36K_^PP^ by the improved counterparts, should they appear, while maintaining the rest of the hereby described HEMSE pipeline.

## METHODS

All detailed methods can be found in the accompanying **Transparent Methods** in the Supplemental Information File

## SUPPLEMENTAL INFORMATION

Supplemental Information can be found in the attached File

## ACKNOWLEDGMENTS

Authors are indebted to Sebastian S. Cocioba for posting in Twitter the useful recipe of LB-charcoal medium (https://twitter.com/ATinyGreenCell/status/1037332606432555009) and to Csaba Pal (Institute of Biochemistry, Biological Research Centre, Szeged) for his continued support. This work was funded by the MADONNA (H2020-FET-OPEN-RIA-2017-1-766975), BioRoboost (H2020-NMBP-BIO-CSA-2018), and SYNBIO4FLAV (H2020-NMBP/0500) Contracts of the European Union and the S2017/BMD-3691 InGEMICS-CM funded by the Comunidad de Madrid (European Structural and Investment Funds).

## AUTHOR CONTRIBUTIONS

TA, EMG, AN and VdL designed the study. TA run the experiments. TA, EMG and VdL wrote the manuscript.

## SUPPLEMENTARY INFORMATION

### TRANSPARENT METHODS

#### Strains and media

The bacterial strains employed in this study are listed in Supplementary Table S2. *E. coli* and *P. putida* strains were grown in liquid LB with shaking (170 rpm) at 37 °C and 30 °C, respectively (Sambrook et al., 1989) with the exception of *E. coli* strains bearing SEVA plasmids endowed with the *c*I857-P*L* thermo-inducible expression system (cargo #14; i.e. pSEVA2514-*rec2*-*mutL*_E36K_^PP^ and derivatives), which were grown at 30 °C to avoid promoter activation. After electroporation recovery during recombineering experiments was performed in Terrific Broth without glycerol (TB: 12 g l^-1^ tryptone, 24 g l^-1^ yeast extract, 2 g l^-1^ KH_2_PO_4_, 9.4 g l^-1^ K_2_HPO_4_). M9 minimal media was prepared according to (Sambrook et al., 1989). Solid media was prepared adding 15 g/L^-1^ of agar to liquid media. M9 solid media was supplemented with 0.2% (w/v) citrate and appropriate antibiotics to select *P. putida* cells in mating experiments. Liquid and solid media were added, when necessary, with 50 μg ml^-1^ of kanamycin (Km), 15 μg ml^-1^ of gentamicin (Gm) for *P. putida* and 10 μg ml^-1^ of the same antibiotic for *E. coli*, 30 μg ml^-1^ of chloramphenicol (Cm), 100 μg ml^-1^ of streptomycin (Sm), 100 μg ml^-1^ of rifampicin (Rif), 50 μg ml^-1^ of nalidixic acid (Nal), 20 μg ml^-1^ of Uracil (Ura), 250 μg ml^-1^ of 5-fluoroorotic acid (5-FOA) and 5 mM of benzoic acid (pH 11). For screening of fluorescent colonies, LB solid media was prepared with 1 mg ml^-1^ of activated charcoal (Sigma-Aldrich Ref. C9157-500G) in order to better discriminate low-signal colonies. Activated charcoal was added to the LB-Agar prior autoclaving and the media poured into 150 mm Petri dishes after vigorous shaking to evenly distribute the insoluble charcoal particles.

#### General procedures, primers and bacterial transformation

Standard DNA manipulations were carried out following routine protocols (Sambrook et al., 1989) and according to manufacturer recommendations. Isothermal Assembly was performed with Gibson Assembly^®^ Master Mix (New England Biolabs, Ipswich, MA, USA). Plasmidic DNA was purified with the QIAprep^®^ Spin Miniprep Kit, both purchased from Qiagen (Valencia, CA, USA). DNA Amplitools Master Mix (Biotools, Madrid, Spain) was used for diagnosis PCRs and amplification of DNA fragments for cloning purposes was done with Q5 polymerase (New England Biolabs, Ipswich, MA, USA). Synthetic oligonucleotides used in this study are listed in Supplementary Table S1 and were purchased from Sigma-Aldrich (St. Louis, MO, USA). PCR products were purified with the Nucleospin^®^ Gel and PCR Clean-up Kit (Macherey-Nagel, Düren, Germany). DNA sequencing was performed in Macrogen (Spain). Transformation of *E. coli* strains was carried out with chemically competent cells using the CaCl_2_ method (Sambrook et al., 1989). Plasmids were introduced in *P. putida* strains via tripartite mating as described in (Martinez-Garcia and de Lorenzo, 2012) and selected in solid M9 minimal media supplemented with 0.2% w/v citrate and appropriate antibiotics. Tetra-parental mating was used as described by (Choi et al., 2005) to insert the mini-transposon Tn7-M-P_*EM7*_-*gfp*-RBS^−^ into the *att*Tn7 site of *P. putida* EM42, using M9-citrate-Gm as selective media (see below for details).

#### Construction of plasmids and strains

The medium-high copy number plasmid pSEVA2514-*rec2*-*mutL*_E36K_^PP^ (Supplementary Table S2) was used for the construction of two derivatives bearing low-and medium-copy number origins of replication. This plasmid was cut with PacI/SpeI and the 4.2 Kb DNA band, containing the *rec2* and *mutL*_E36K_^PP^ genes under the control of the thermo-inducible system *c*I857/P_L_, was ligated to PacI/SpeI restricted plasmids pSEVA221 (low copy number) and pSEVA231 (medium copy number). Ligations were transformed into *E. coli* CC118 and selection was made in LB-Km plates, obtaining plasmids pSEVA2214-*rec2*-*mutL*_E36K_^PP^ and pSEVA2314-*rec2*-*mutL*_E36K_^PP^. Both constructs were separately introduced in *P. putida* EM42 by tri-parental matings followed by selection in M9-citrate-Km solid media, obtaining the strains *P. putida* EM42 (pSEVA2214-*rec2*-*mutL*_E36K_^PP^) and *P. putida* EM42 (pSEVA2314-*rec2*-*mutL*_E36K_^PP^). *P. putida* EM42 was also transformed by the same method with pSEVA2314, generating the control strain *P. putida* EM42 (pSEVA2314). A Tn*7* mini-transposon with the *gfp* gene under the control of the constitutive P_*EM7*_ promoter, but lacking the ribosome binding site (RBS) sequence, was constructed. To this end, first the *gfp* gene was placed under the control of the P_*EM7*_ promoter: plasmid pSEVA637 (Supplementary Table S2) was cut with HindIII/SpeI and the purified 0.7 Kb band (RBS + *gfp* gene) was ligated to the pSEVA237R-PEM7 (Supplementary Table S2) backbone digested with HindIII/SpeI. Upon transformation in *E. coli* CC118 and selection on LB-Km plates, the resulting plasmid (pSEVA237-PEM7) was digested with PacI/SpeI and the purified 0.9 Kb band (containing the *gfp* gene under the control of the P_*EM7*_ and bearing a consensus 5’-AGGAGG-3’ RBS sequence) was ligated to a pTn7-M plasmid restricted with the same enzymes. Ligation mixture was used to transform *E. coli* competent cells and selection was done in LB-KmGm plates. The resulting plasmid, pTn7-M-PEM7-GFP, was used as a template to eliminate the RBS sequence. In order to achieve this, the plasmid was PCR amplified with primers Tn7-PEM7-F/ Tn7-PEM7-R (Tm= 58 °C, 2 min. elongation, Q5 polymerase). The primers were designed to i) amplify the whole plasmid with the exception of the 7-nt Shine Dalgarno motif 5’-AGGAGGA-3’ located 7-nt away from the *gfp* start codon, ii) generate an PCR product sharing a 40-bp sequence at both sides of the molecule to allow isothermal assembly of the amplicon. The 3.9 Kb PCR product was purified and subjected to Gibson Assembly and the reaction was transformed into *E. coli*. Selection was made in LB-KmGm plates, thus obtaining the plasmid pTn7-M-P_*EM7*_-GFP-RBS^−^. The region between the P_*EM7*_ promoter and the end of the *gfp* gene was fully sequenced with primers PS2 and PEM7-F to ensure the correct deletion of the RBS sequence. *E. coli* (pTn7-M-P_*EM7*_-GFP-RBS^−^) was used as the donor strain to introduce the mini-transposon in the *att*Tn7 site of *P. putida* EM42. Both strains and the helper strains *E. coli* HB101 (pRK600) and *E. coli* (pTNS2) were used in a tetra-parental mating followed by selection in M9-citrate-Gm solid media. Colonies were streaked in the same media and subjected to two diagnostic PCRs to check the mini-transposon insertion. PCRs with primer pairs PS2/ PP5408-F (Tm= 60 °C, 1 min. 30 seconds elongation) and PEM7-F/Tn7-GlmS (Tm= 60 °C, 1 min. elongation) yielded bands of 2.2 Kb and 1.2 Kb, respectively, confirming the correct integration of the transposon in the *att*Tn7 locus. The resulting strain *P. putida* EM42::Tn*7*-M-P_*EM7*_-*gfp*-RBS^−^ (referred as *P. putida* TA245 in Supplementary Table S2) was transformed by tripartite mating with pSEVA2314-*rec2*-*mutL*_E36K_^PP^ plasmid. After selection on M9-citrate-KmGm plates, the strain *P. putida* TA245 (pSEVA2314-*rec2*-*mutL*_E36K_^PP^) was obtained. Integrity of the constructs described above, either in *E. coli* or *P. putida*, was always checked by miniprep, restriction and agarose gel visualization.

#### Oligonucleotide design, recombineering protocol, cycling procedure and screening

The nine oligonucleotides used in this work for recombineering experiments (SR, NR, RR, PR, CR, RBS-C_6_, RBS-Deg_6_, RBS-C_9_, RBS-Deg_9_) were designed to introduce different allelic changes targeting the lagging strand of the *P. putida* chromosome. Supplementary Table S3 summarizes the main features of each oligonucleotide while complete sequence and additional details can be found in Supplementary Table S1. The recombineering protocol used here relies in the coexpression of the Rec2 recombinase and the MutL_E36K_^PP^ dominant negative allele from plasmids endowed with the thermo-inducible *c*I857/P_L_ expression system (pSEVA2214-*rec2*-*mutL*_E36K_^PP^, pSEVA2314-*rec2*-*mutL*_E36K_^PP^ or pSEVA2514-*rec2*-*mutL*_E36K_^PP)^. The protocol is basically identical to that described previously in (Aparicio et al., 2019b). Overnight cultures of *P. putida* strains harboring the proper plasmid were used to inoculate 20 ml of fresh LB-Km at OD_600_ = 0.1 in 100 ml Erlenmeyer flasks. Cultures were incubated at 30 °C with vigorous shaking (170 rpm) until OD_600_ ~ 1.0 and flasks were then placed in a water bath at 42 °C for 5 minutes to increase rapidly the temperature and induce the P_*L*_ promoter. Ten additional minutes of incubation at 42 °C was performed in an air shaker at 250 rpm (induction total time at 42 °C= 15 minutes) to trigger the expression of *rec2*-*mutL*_E36K_^PP^ genes, followed by 5 minutes in ice to cool down the bacterial culture and stop the induction. Competent cells were then prepared transferring 10 ml of each culture to 50-ml conical tubes and centrifuging the cells at 3,220 g/ 5 minutes. Cell pellets were resuspended in 10 ml of 300 mM sucrose and washed two additional times with 5 and 1 ml of the same solution. After centrifugation in a bench-top centrifuge (10,000 rpm, 1 minute), cellular pellets were finally resuspended in 200 μl of 300 mM sucrose and 100 μl of this suspension was added with the recombineering oligonucleotide. For single-oligonucleotide experiments, 1 μl from a 100 μM stock was used (1 μM final concentration). For multiplexed experiments, 10 μl of each oligonucleotide stock at 100 μM (SR, NR, RR, PR and CR) were mixed and 3 μl of this mixture were added to the competent cells (accounting for 0.6 μM of each oligo.). The cell suspension was mixed thoroughly by pipetting, placed in an electroporation cuvette (Bio-Rad, 2 mm-gap width) and electroporated at 2.5 kV in a Micropulser™ device (Bio-Rad Laboratories, Hercules, CA, USA). Cells were immediately inoculated in 5 ml of fresh TB in 100 ml Erlenmeyer flaks and recovered at 30 °C/ 170 rpm. Before plating the cells for screening of allelic replacements, different recovery times and TB additions were used depending on the experiment. For one cycle recombineering experiments, overnight recovery was done in TB for assays with SR and NR oligonucleotides while for experiments with oligonucleotides RBS-C_6_, RBS-Deg_6_, RBS-C_9_ and RBS-Deg_9_, TB supplemented with Km and Gm was preferred. Specifications for cycled recombineering assays (HEMSE) are depicted below.

#### High-efficiency multi-site genomic editing protocol

HEMSE is a cycled recombineering protocol run in a multiplexed fashion. The procedure involves a standard recombineering protocol in which, as explained before, cultures were subjected to electrotransformation with an equimolar mixture of several oligonucleotides. The recovery was performed in TB added with Km in order to maintain the plasmid along the cycles, and the incubation proceeded at 30 °C with vigorous shaking (170 rpm) until an OD_600_ ~ 1.0 (Cycle-I). Culture aliquots were withdrawn for screening and the bacterial culture entered in the next round of recombineering by performing induction at 42 °C/ 15 minutes, competent cell preparation, oligonucleotide mixture electroporation and recovery till reaching again a cell density around 1.0 at 600 nm (Cycle-II). Further cycles proceeded in the same way (Fig. 3 of main text). Each cycle took one day in average and recovery, when necessary, was performed overnight at room temperature without shaking to avoid culture overgrowth. When recovery step was completed at the end of the day, cultures were stored at 4 °C overnight. A new cycle was started in the next morning incubating the culture 30 minutes at 30 °C (170 rpm) before the induction step. Screening of allelic changes after recombineering was performed plating aliquots of recovered cultures in the appropriate selective and/or non-selective solid media, as follows:

- In single-olignucleotide experiments with SR and NR oligonucleotides (one cycle), overnight cultures were plated in LB-Sm (dilutions 10^-4^ and 10^-5^) and LB-Nal (dilutions 10^-4^ and 10^-5^), respectively, to estimate the allelic replacements, while dilutions 10^-7^ and 10^-8^ were done in LB without antibiotics to count viable cells. Plates were incubated 18 h. at 30 °C and CFUs annotated.
- In single-oligonucleotide experiments with RBS-C_6_, RBS-Deg_6_, RBS-C_9_, RBS-Deg_9_ oligos (one cycle), cultures recovered overnight were plated on 150 mm width LB-KmGm-activated charcoal plates using 10^-6^ dilutions. This allowed an average of 500 colonies per plate. To facilitate the identification of colonies displaying low levels of fluorescence, plates were incubated at 30 °C for 5 days. Fluorescent colonies were streaked in the same media and insertion of putative ribosome binding sites upstream the *gfp* gene were checked by PCR amplifying this DNA region with primers PS2/ PP5408-F (Tm= 60 °C, 1 min. 30 seconds elongation, 1.0 Kb product) and sequencing the amplicon with primer ME-I-Gm-ExtR. Non-redundant clones with different sequences inserted were selected and glycerol stocks made prior characterization by flow cytometry.
- Allelic replacements in HEMSE experiments were screened after recovery steps (OD_600_ ~ 1.0) of cycle-I, cycle-V and cycle-X. Viable cells were estimated plating dilutions 10^-7^ and 10^-8^ in LB plates. Single mutants coming from SR-, NR-, RR- and PR-mediated recombineering were analyzed by plating dilutions 10^-4^ and 10^-5^ in LB-Sm, LB-Nal, LB-Rif and LB-5FOA-Ura plates. Plates were incubated 24 h at 30 °C and total CFUs of single mutants (Sm^R^, Nal^R^, Rif^R^ and 5FOA^R^) and viable cells were taken. Twenty 5FOA^R^ colonies were replicated on M9-citrate and M9-citrate-5FOA-Ura plates in order to discriminate authentic *pyrF* mutants (5FOA^R^/Ura^-^) from spontaneous 5FOA^R^ mutants (5FOA^R^/Ura^+^), as stated in (Galvao and de Lorenzo, 2005; Aparicio et al., 2016). Colonies grown on both media were discounted of the total 5FOA^R^ numbers as *pyrF*-unrelated, spontaneous mutants. Dilutions 10^-6^ in LB-benzoate plates allowed the estimation of *catA*-I^-^ mutants simply by counting the dark-brown colonies appeared after 10 days of incubation at 30 °C. *catA*-I^-^ mutants accumulate catechol, which turns into brown intermediates after spontaneous oxidation and polymerization (Jimenez et al., 2014). In previous assays aimed to obtain *catA*-I^-^ mutants through recombineering with CR oligo, it was noticed that long incubations were necessary to appreciate the colored phenotype in solid media (Fig. S2). The observed dark-brown colonies were always *catA*-I^-^ mutants, as was demonstrated by amplification of *catA*-I gene (primers catA-F/catA-R, Tm 55 °C, 1 minute elongation) and sequencing of the 0.5 Kb amplicon with primer catA-F (data not shown) in 20 selected colonies. Multiple gene editions were also analyzed plating cultures from cycles I, V and X on LB solid media supplemented either with Sm+Nal+ Rif+5FOA+Ura (four editions mediated by SR, NR, RR and PR oligonucleotides), 24 incubation at 30 °C, or with Sm+Nal+Rif+5FOA+Ura+benzoate (five editions mediated by the 5 oligonucleotides used in this study), 10 days incubation at 30 °C. For this last experiment, there were considered quintuple mutants those colonies displaying resistance to Sm, Nal, Rif and 5FOA and also showing the characteristic brown phenotype of *catA*-I^-^ mutants. The recombineering frequency (RF) was calculated as the ratio between the number of colonies showing a given phenotype and the number of viable cells within the experiment, being this ratio normalized to 10^9^ viable cells for graphic representation.

In order to check the accuracy of the allelic replacements, 18 colonies showing the quintuple mutant phenotype (Sm^R^, Nal^R^, Rif^R^, 5FOA^R^ and catechol accumulation) were checked by PCR amplification and sequencing of the target genes. For each bacterial clone, five different PCRs were set up to amplify: *rpsL* (primers rpsL-Fw/ rpsL-Rv, Tm 57 °C, 45 seconds elongation, 0.8 Kb product), *gyrA* (primers gyrA-Fw/ gyrA-Rv, Tm 57 °C, 45 seconds elongation, 0.4 Kb product), *rpoB* (primers rpoB-F/rpoB-R, Tm 57 °C, 45 seconds elongation, 0.4 Kb product), *pyrF* (primers pyrF-F/pyrF-R, Tm 52 °C, 1 minute elongation, 1.2 Kb product) and *catA-I* (primers catA-F/catA-R, Tm 55 °C, 1 minute elongation, 0.5 Kb product). The purified PCR products were sequenced with the putative forward primers and the sequence analysed for the expected changes mediated by recombineering. All clones analysed (n=18; 100%) showed the correct changes, demonstrating that the observed phenotypes corresponded to mutations mediated by the HEMSE procedure. Single allelic replacements in HEMSE experiments were not confirmed by PCR and sequencing since previous works showed that virtually 100 % of Sm^R^, Nal^R^ and *pyrF*^-^ mutants obtained by recombineering with oligos SR, NR and LM (almost identical to RR oligo used in this work) harbored the expected changes in the target genes *rpsL*, *gyrA* and *pyrF* (Ricaurte et al., 2018) (Aparicio et al., 2019a). Preliminary studies in this work showed, on the other hand, that single Rif^R^ mutants also displayed 100 % accuracy in the expected mutations of the target gene *rpoB* (data not shown). As explained before, *catA*-I^-^, dark-brown colonies were also analyzed by PCR and sequencing in previous test experiments (data not shown), with analogous results.

#### Flow cytometry

The visual selection of fluorescent colonies from recombineering experiments with oligos RBS-Deg_6_ and RBS-Deg_9_ gave rise to a collection of 31 RBS insertion mutants showing a wide variety of fluorescent signals. Together with the negative controls of *P. putida* TA245 (insertion of Tn*7*-M-P_*EM7*_-*gfp*-RBS^−^) and *P. putida* EM42 (no *gfp* gene), a total of 33 strains were characterized for GFP production. Each strain was inoculated from glycerol stocks in 2 ml of LB-KmGm (*P. putida* EM42 in LB; *P. putida* TA245 in LB-Gm) and cultured at 30 °C/ 170 rpm. 0.5 ml of overnight cultures (OD_600_ ~ 2-3) were centrifuged and resuspended in 1 ml of filtered Phosphate Buffered Saline (PBS) 1X (8mM Na2HPO4, 1.5mM KH2PO4, 3mM KCl, 137mM NaCl, pH.7.0). Fifty μl of each suspension was added to 450 μl of PBS 1X to obtain cellular samples with OD_600_ ~ 0.10.15. Samples were analyzed in a MACSQuant™ VYB cytometer (Miltenyi Biotec, Bergisch Gladbach, Germany) to quantify the emission of fluorescence as indicated in (Martinez-Garcia et al., 2014). GFP was excited at 488 nm and the fluorescence signal was recovered with a 525±40 nm band-pass filter. For each sample, at least 100,000 events were analyzed and the FlowJo v. 9.6.2 software (FlowJo LLC, Ashland, OR, USA) was used to process the results. Population was gated to eliminate background noise and the median of the GFP-A channel of two biological replicas was used for graphical representation.

## SUPPLEMENTARY FIGURES

**Figure S1.**
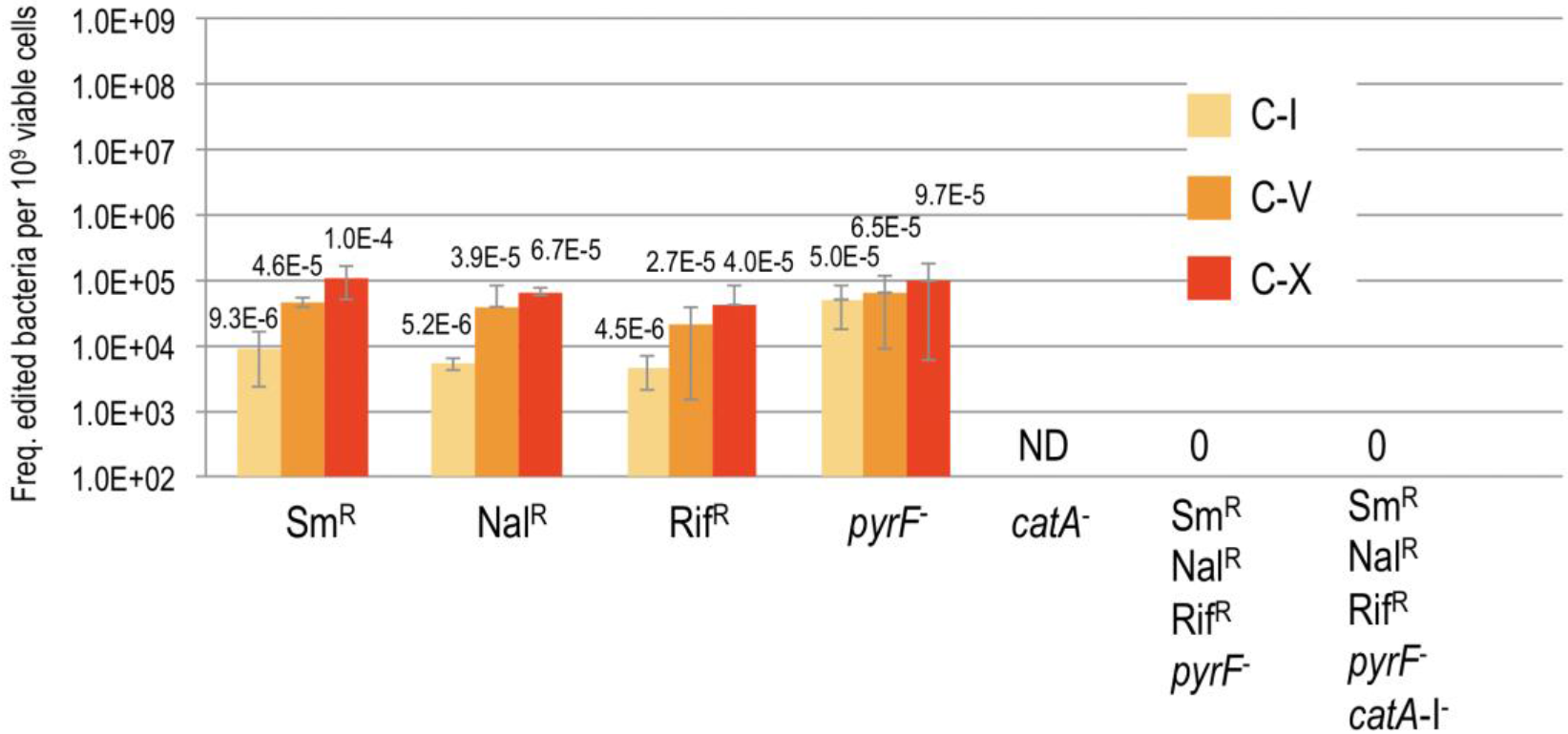
Rec2-independent editing in HEMSE assays. Editing efficiencies of single and multiple changes in the control strain *P. putida* EM42 harboring the empty plasmid pSEVA2314 were assayed applying 10 cycles of HEMSE and an equimolar mixture of oligos SR, NR, RR, PR and CR, following the same procedure explained in Figure 4A and 4B. See more details in Transparent Methods section. Allelic replacements of *catA*-I gene were not determined in these assays (ND), while multiple editions could not be detected (0).

**Figure S2.**
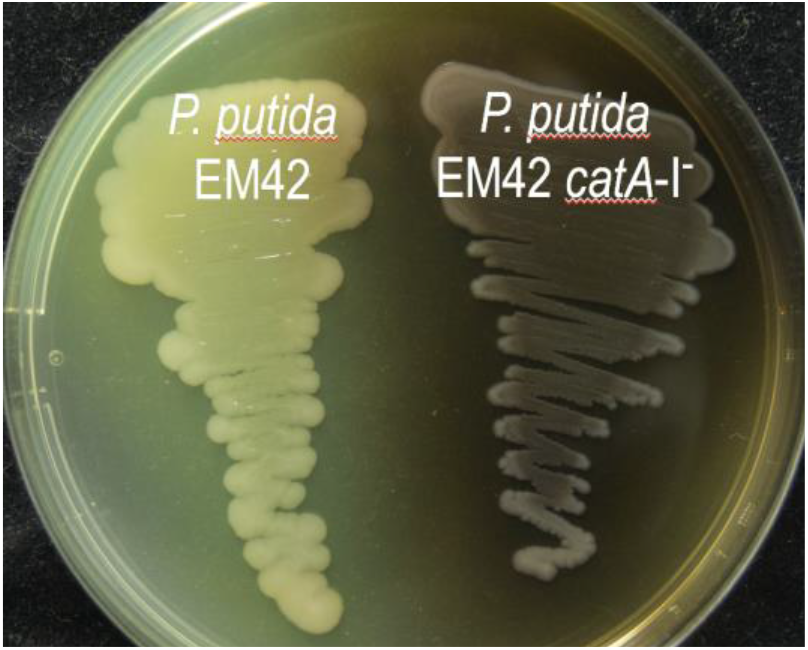
Phenotype of *P. putida* EM42 *catA*-I^-^ strain. *P. putida* EM42 (pSEVA2314-*rec2*-*mutL*_E36K_^PP^) was subjected to recombineering with CR oligonucleotide (see Transparent Methods section). Three stop codons were inserted in the *catA*-I ORF, generating a mutant strain in which the metabolism of benzoic acid is impaired, leading to accumulation of catechol (Jimenez et al., 2014). Upon spontaneous oxidation and polymerization, catechol derivatives exhibit a characteristic dark-brown colour. *P. putida* EM42 and the catA-I^-^ mutant were grown in LB-Agar supplemented with benzoate 5 mM and incubated 10 days at 30 °C to allow the visualization of the colored phenotype.

## SUPPLEMENTARY TABLES

**SUPPLEMENTARY TABLE S1.**
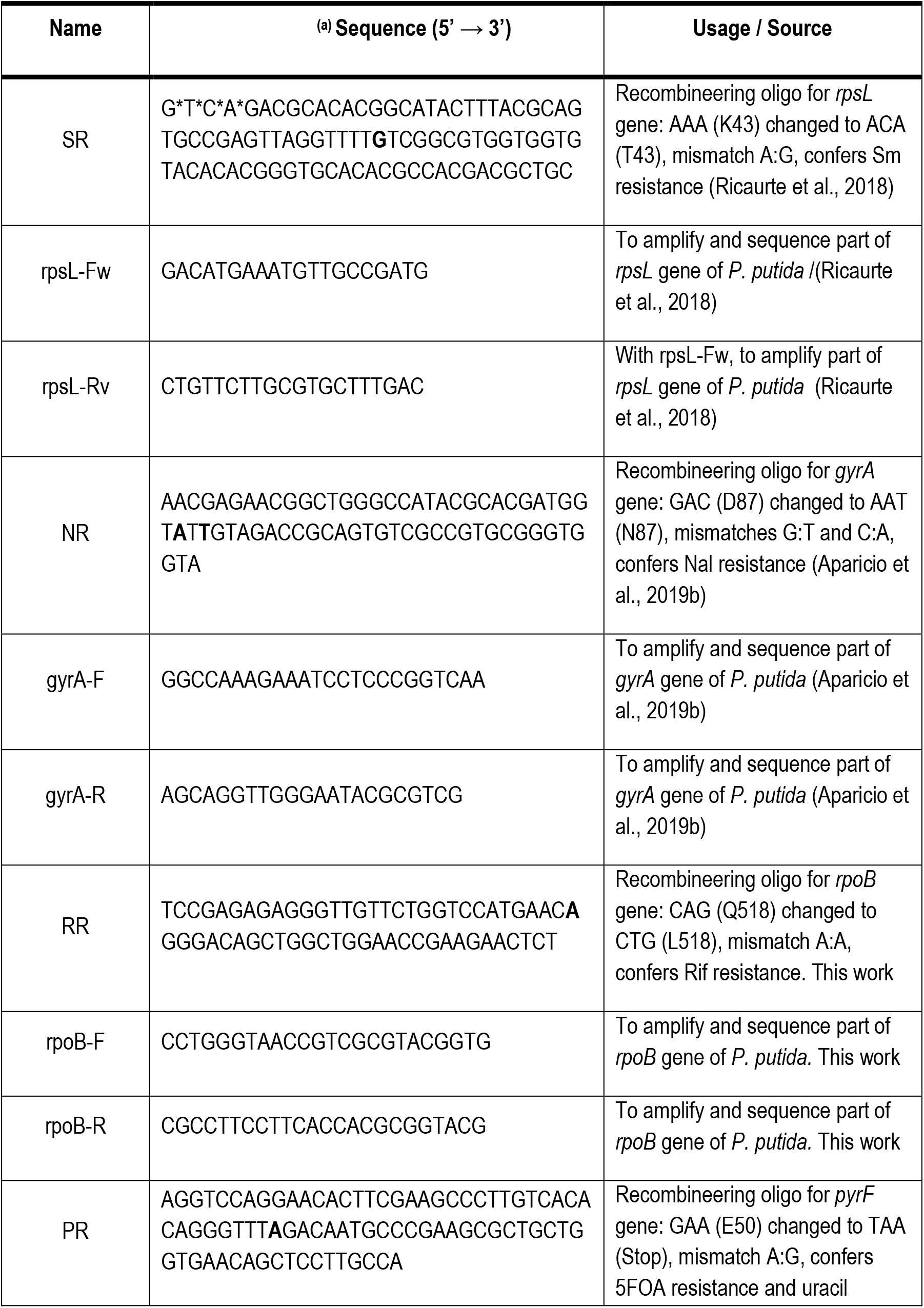
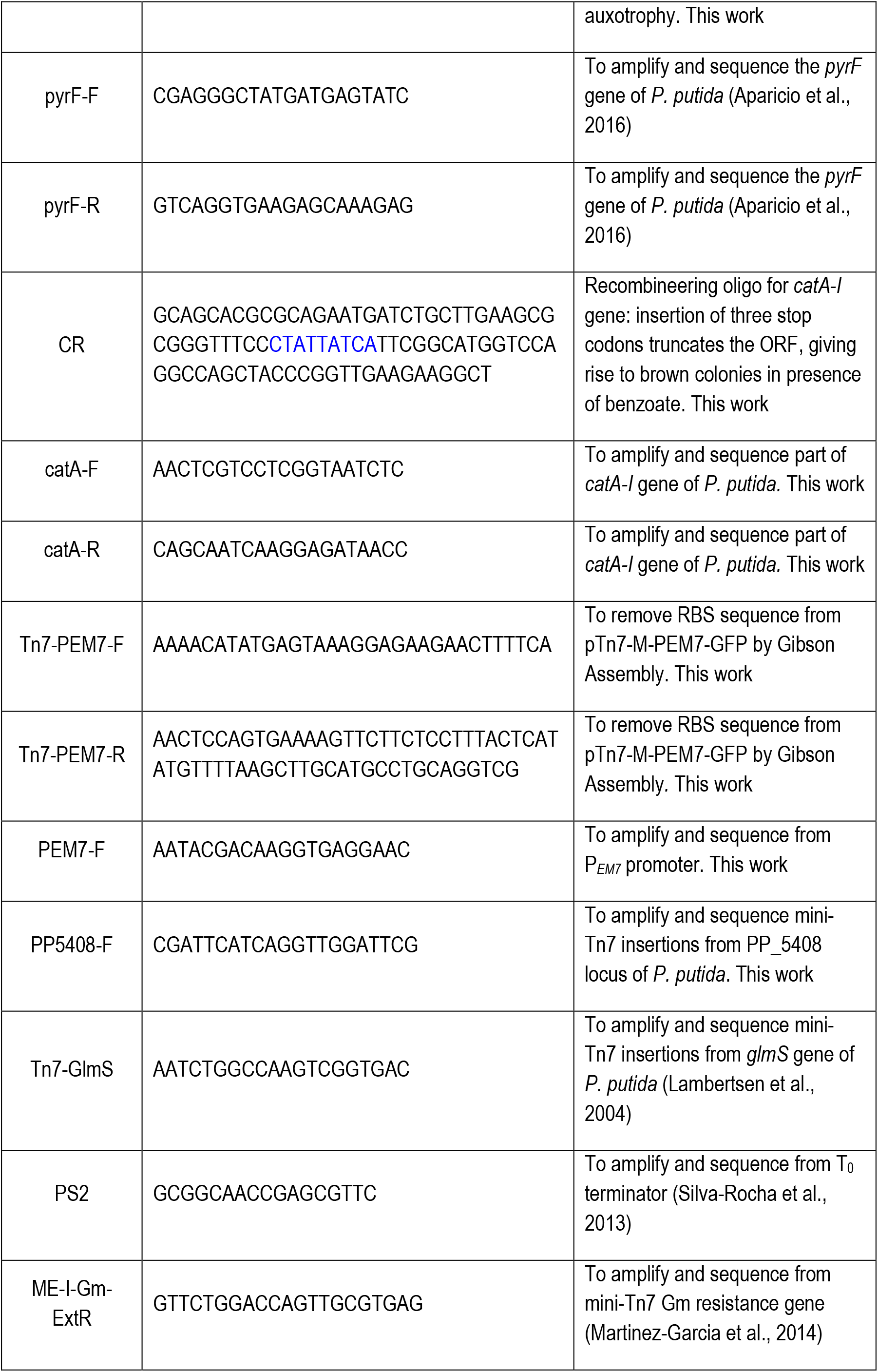
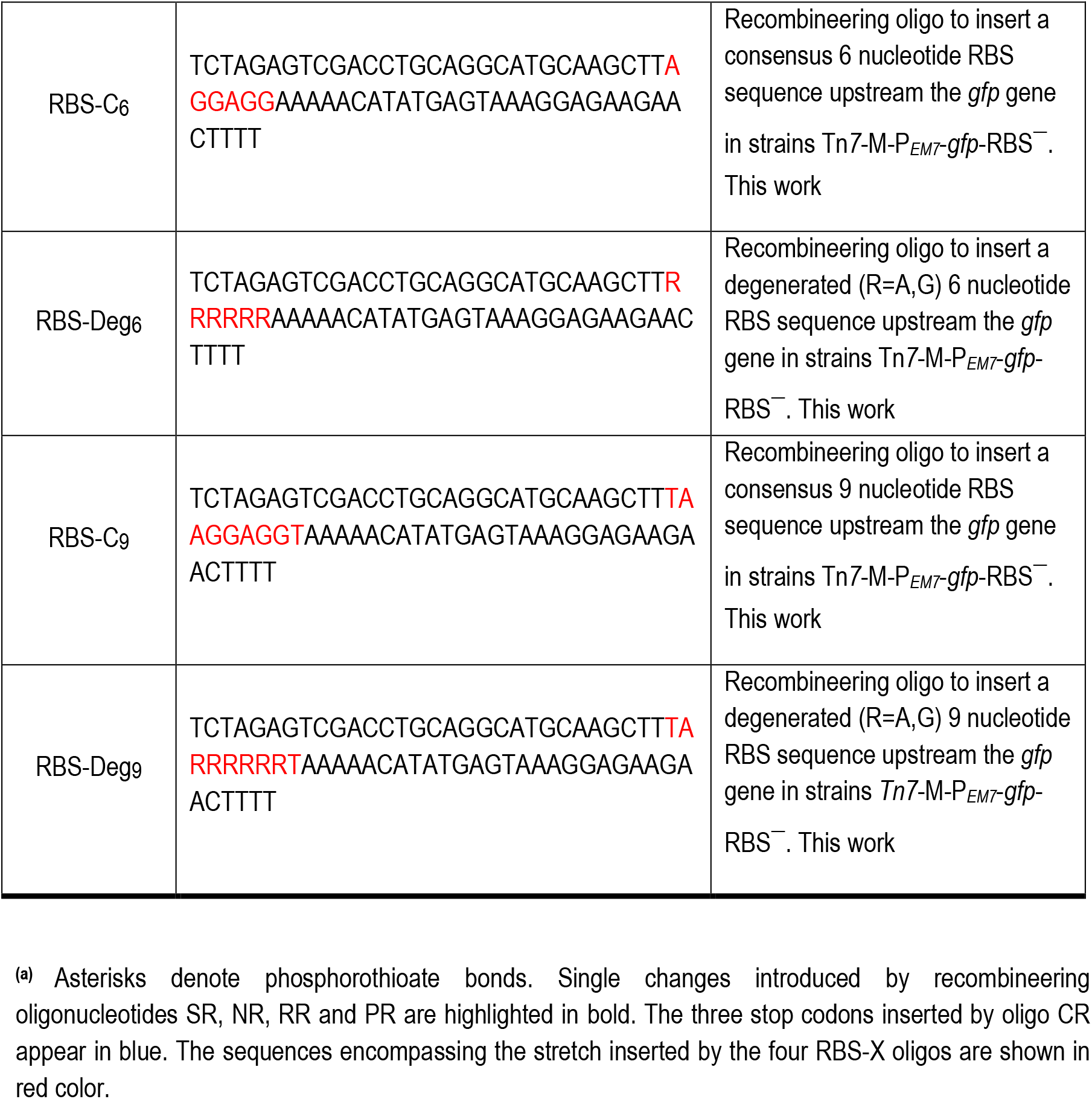
Oligonucleotides used in this study.

**SUPPLEMENTARY TABLE S2.** Bacterial strains and plasmids used in this work.

| Strain or plasmid | Relevant characteristics <sup>a</sup> | Reference or source |
| --- | --- | --- |
| *Escherichia coli* |  |  |
| CC118 | Cloning host; $\Delta(ara-leu)$ *araD* $\Delta lacX174$ *galE* *galK* *phoA* *thiE1* *rpsE*(Sp<sup>R</sup>) *rpoB*(Rif<sup>R</sup>) *argE*(Am) *recA1* | (Manoil and Beckwith, 1985) |
| HB101 | Helper strain used for conjugation; F<sup>-</sup> $\lambda^-$ *hsdS20*(rB<sup>-</sup> mB<sup>-</sup>) *recA13* *leuB6*(Am) *araC14* $\Delta(gpt-proA)62$ *lacY1* *galK2*(Oc) *xyl-5* *mtl-1* *thiE1* *rpsL20*(Sm<sup>R</sup>) *glnX44* (AS) | (Boyer and Roulland-Dussoix, 1969) |
| CC118λpir | Cloning host for plasmids containing an R6K origin of replication; $\Delta(ara-leu)$ *araD* $\Delta lac$ X174 *galE* *galK* *phoA20* *thi-1* *rpsE* *rpoB* *argE* (Am) *recA1*, λpir lysogen | (Herrero et al., 1990) |
| *Pseudomonas putida* |  |  |
| EM42 | KT2440 derivative; Δprophage1 Δprophage4 Δprophage3 Δprophage2 ΔTn7 ΔendA-1 ΔendA-2 ΔhsdRMS Δflagellum ΔTn4652 | (Martinez-Garcia et al., 2014) |
| TA238 | EM42 derivative; *rpsL*<sup>-</sup>(Sm<sup>R</sup>) *gyrA*<sup>-</sup>(Nal<sup>R</sup>) *rpoB*<sup>-</sup>(Rif<sup>R</sup>) *pyrF*<sup>-</sup>(5FOA<sup>R</sup>) *catA-I*<sup>-</sup> | This work |
| TA245 | EM42 derivative with mini-Tn7-M-P<sub>EM7</sub>-gfp-RBS<sup>-</sup> transposon inserted in the attTn7 site | This work |
| Plasmids |  |  |
| pSEVA2314 | Inducible expression vector; *oriV*(pBBR1); cargo [cl857-P<sub>L</sub>]; standard multiple cloning site; Km<sup>R</sup> | (Aparicio et al., 2019a) |
| pSEVA2214-*rec2*-*mutL*<sub>E36K</sub><sup>PP</sup> | pSEVA2214 derivative bearing the *rec2* recombinase and *mutL*<sub>E36K</sub><sup>PP</sup> allele ; *oriV*(RK2); cargo [ cl857-P<sub>L</sub> → *rec2*-*mutL*<sub>E36K</sub><sup>PP</sup>]; Km<sup>R</sup> | This work<br>GenBank n° MN688223 |
| pSEVA2314-*rec2*- *mutL*<sub>E36K</sub><sup>PP</sup> | pSEVA2314 derivative bearing the *rec2* recombinase and | This work |
|  | *mutL<sub>E36K</sub><sup>PP</sup>* allele ; *oriV*(pBBR1); cargo [ *cl857-P<sub>L</sub>* → *rec2-mutL<sub>E36K</sub><sup>PP</sup>*]; Km<sup>R</sup> | GenBank n°<br>MN688222 |
| pSEVA2514-*rec2-mutL<sub>E36K</sub><sup>PP</sup>* | pSEVA2514 derivative bearing the *rec2* recombinase and *mutL<sub>E36K</sub><sup>PP</sup>* allele ; *oriV*(RFS1010); cargo [ *cl857-P<sub>L</sub>* → *rec2-mutL<sub>E36K</sub><sup>PP</sup>*]; Km<sup>R</sup> | (Aparicio et al.,<br>2019b)<br>GenBank n°<br>MN180222 |
| pSEVA637 | *oriV*(pBBR1); cargo [*gfp*]; Gm<sup>R</sup> | (Silva-Rocha et<br>al., 2013)<br>(Martinez-Garcia et al.,<br>2015) |
| pSEVA237R-PEM7 | *oriV*(pBBR1); cargo [ *P<sub>EM7</sub>* → *mCherry*]; Km<sup>R</sup> | (Silva-Rocha et<br>al., 2013)<br>(Martinez-Garcia et al.,<br>2015) |
| pSEVA237-PEM7 | *oriV*(pBBR1); cargo [ *P<sub>EM7</sub>* → *gfp*]; Km<sup>R</sup> | This work |
| pTn7-M | *oriV*(R6K); mini-Tn7 transposon; standard multiple cloning site; Km<sup>R</sup> Gm<sup>R</sup> | (Zobel et al.,<br>2015) |
| pTn7-M-PEM7-GFP | pTn7-M derivative with *P<sub>EM7</sub>-gfp* in the mini-Tn7 transposon; Km<sup>R</sup> Gm<sup>R</sup> | This work |
| pTn7-M-PEM7-GFP-RBS<sup>-</sup> | pTn7-M-PEM7-GFP derivative lacking the *gfp* RBS; Km<sup>R</sup> Gm<sup>R</sup> | This work |
| pRK600 | Helper plasmid used for conjugation; *oriV*(ColE1), RK2 (mob+ tra+); Cm<sup>R</sup> | (Kessler et al.,<br>1992) |
| pTNS2 | Helper plasmid for mini-Tn7 transposition; *oriV*(R6K), TnsABC+D specific transposition pathway; Ap<sup>R</sup> | (Choi et al.,<br>2005) |

<sup>a</sup> Antibiotic markers: Ap, ampicillin; Cm, chloramphenicol; Km, kanamycin; 5FOA, 5-fluoro-orotic acid;
Gm, gentamicin; Nal, nalidixic acid; Rif, rifampicin; Sm, streptomycin; Sp, spectinomycin.

**SUPPLEMENTARY TABLE S3.** Main features of recombineering oligonucleotides used in this study.

| Name | P-thioate bonds | Length | Target gene | Change/mismatch | MMR sensitivity | $\Delta G$ (kcal/mol) | Phenotype |
| --- | --- | --- | --- | --- | --- | --- | --- |
| SR | Four at 5'-end | 94 | *rpsL* | A→C/A:G | Low | - 20.79 | Sm<sup>R</sup> |
| NR | None | 65 | *gyrA* | G→A/ G:T<br>C→T/ C:A | High | - 11.84 | Nal<sup>R</sup> |
| RR | None | 60 | *rpoB* | A→T/ A:A | High | - 7.26 | Rif<sup>R</sup> |
| PR | None | 81 | *pyrF* | G→T/ G:A | Low | - 14.03 | 5-FOA<sup>R</sup>/Ura<sup>-</sup> |
| CR | None | 89 | *catA-I* | Insertion 3 Stops | Low | - 12.84 | Catechol accumulation (Brown color) |
| RBS-C<sub>6</sub> | None | 67 | *gfp* UTR | Insertion 7 nt | Low | - 4.94 | Fluorescent |
| RBS-Deg<sub>6</sub> | None | 67 | *gfp* UTR | Insertion 7 nt (deg.) | Low | Variable | Fluorescent |
| RBS-C<sub>9</sub> | None | 70 | *gfp* UTR | Insertion 10 nt | Low | - 4.59 | Fluorescent |
| RBS-Deg<sub>9</sub> | None | 70 | *gfp* UTR | Insertion 10 nt (deg.) | Low | Variable | Fluorescent |

